# A family of linear plasmid phages that detect a quorum-sensing autoinducer exists in multiple bacterial species

**DOI:** 10.1101/2025.07.30.667625

**Authors:** Francis J. Santoriello, Bonnie L. Bassler

## Abstract

Temperate phages oscillate between lysogeny, a genomic maintenance state within a bacterial host, and lytic replication, in which the host is killed, and newly made phage particles are released. Successful transmission to new hosts requires that temperate phages appropriately time their transitions from lysogeny to lysis. It is well understood that temperate phages trigger lysis upon detection of host cell stress. Understanding of the breadth of cues that induce lysis expanded with the discovery of phages carrying quorum-sensing receptor genes that promote lytic induction exclusively at high host cell density. Bacteria engage in a cell-cell communication process called quorum sensing, which relies on the production, release, accumulation, and group-wide detection of extracellular signal molecules called autoinducers. Bacteria use quorum sensing to monitor changes in population density and synchronize collective behaviors. The temperate phage VP882 (φVP882) encodes VqmAφ – a homolog of its host’s quorum-sensing receptor/transcription factor VqmA. VqmAφ allows φVP882 to detect the accumulation of the host autoinducer called DPO. Presumably, launching the lytic induction program at high host cell density maximizes φVP882 transmission to new hosts. Here, by mining sequence databases for linear plasmid phages, we identify VP882-like phages in multiple DPO-producing bacterial species isolated at diverse times and geographic locations. We show that the VqmAφ homologs can indeed detect DPO and, in response, activate the lytic pathway. Our observation indicates that φVP882 is a member of a large family of globally-dispersed quorum-sensing-responsive temperate phages.

**IMPORTANCE:** The discovery of quorum-sensing responsive linear plasmid phages has transformed understanding of phage-bacterial interactions by demonstrating inter-domain chemical communication. To date, however, examples of quorum-sensing responsive phages have been sparse. The founding example of such a phage, φVP882, detects a chemical communication signal molecule called DPO that is produced by diverse bacterial species. We investigated whether a family of VP882-like phages might exist that detect and respond to DPO. We find that indeed, VP882-like phages reside in DPO-producing bacterial species isolated at different times and geographic locations, suggesting their wide circulation in the environment. This observation strengthens the evidence for the generality of phage-bacterial inter-domain chemical communication.

## OBSERVATION

The *Vibrio parahaemolyticus* temperate phage VP882 (φVP882) monitors host quorum-sensing-mediated communication using a phage-encoded homolog of the vibrio-specific quorum-sensing receptor VqmA (VqmAφ) (1). At high cell density, the VqmAφ receptor binds a host-produced autoinducer, DPO, triggering production of a phage-encoded antirepressor, Qtip. Qtip sequesters and inactivates the phage cI lysis repressor to initiate lytic replication (1, 2). We hypothesize that this host-DNA-damage-independent pathway to lytic induction allows φVP882 to coordinate lysis with a high vicinal density of potential new host cells, thus maximizing transmission. DPO production depends on threonine dehydrogenase (Tdh) (3), which is highly conserved across bacterial species (NCBI HMM: TIGR00692.1). While host VqmA is restricted to the vibrio genus (1), phage-borne, DPO-dependent VqmAφ-Qtip systems could allow VP882-like phages harbored by any DPO-producing bacterial species to respond to host cell density. Despite this potential for broad pertinency, VP882-like phages initially appeared to be rare in sequence databases. φVP882 was first identified in *V. parahaemolyticus* strain 882 (4). Over a decade later, two VP882- like phages were identified, one in a different *V. parahaemolyticus* strain and one in a *Salmonella enterica* strain (2). Efforts to catalog bacteriophages in various environments have expanded sequence databases (5–9). Considering the breadth of DPO production among bacteria, we searched these enlarged sequence repositories for additional VP882-like phages carrying *vqmA*φ*-qtip*.

We used TelN, a conserved component of the linear plasmid phage replication machinery (10), to query NCBI and 5 phage-specific databases (5–9). We extracted 8,537 unique phage sequences (Supplementary Dataset 1). We clustered the identified phages with the Prokaryotic Viral RefSeq database to identify their genus-level relationships with annotated RefSeq viral sequences. The analysis returned 838 viral clusters (Fig. 1A, Supplementary Dataset 2). We focused on φVP882, which clustered into VC_18_0 with 19 other phage genomes (Fig. 1A, Table S1, Supplementary Dataset 2). This cluster returned a Genus Confidence Score of 1.0, indicating that all phages in the cluster are likely of the genus Hapunavirus, which prior to our analysis, only included φVP882 and φHAP-1 of *Vreelandella aquamarina* (11). Our analysis shows that, in fact, φHAP-1 clusters into VC_20_0 along with 18 other phage genomes (Fig. 1A), indicating that φHAP-1-like Hapunaviruses are distinct from VP882-like Hapunaviruses (Fig. S1). We aligned the genomes of the 20 VP882-like phages and identified conservation over their entire genomes apart from two variable regions (Fig. 1B): one in the structural gene region that encodes a predicted P1-like GpU (UniProt: Q71TD6 · U1_BPP1), likely involved in host tropism, and one covering the accessory region, a common site of genomic innovation (12). To the 20 clustered VP882-like phages, we added two more putative VP882-like phage genomes, harbored by *V. parahaemolyticus* E4_10 and *Shewanella algae* CLS1, that were excluded from our search due to their fragmented natures. The *vqmA*φ*-qtip* module genes are conserved in 17 of the 22 VP882-like phages (Fig. 2A,B). We note that one of the 17 VP882-like phages harbors the *vqmA*φ*-qtip* module at the end of its contig, and thus it cannot be verified as intact (Fig. 2A,B). The 5 phages lacking the module possess fragments of the *vqmA*φ and *qtip* genes, indicating elimination of a portion of the module, presumably via deletion (Fig. 2A,B). Among the 16 VP882-like phages with a complete *vqmA*φ*- qtip* module, variations have yielded 9 VqmAφ isoforms that do not cluster by host species (denoted VqmAφ^1-9^ in Fig. S2A). The Qtip proteins are identical in 14 of the 16 genomes (Fig. S2B). The partner cI proteins to the 14 identical Qtip proteins differ from the φVP882 cI protein by 0 to 11 amino acid substitutions and insertions (Fig. S2C). This finding indicates that the 14 identical Qtip proteins can tolerate these variations. The 2 non-identical Qtip proteins co-occur with more distant cI proteins (68% and 44% amino acid identity to the φVP882 cI; Fig. S2B,C).

**Fig 1.**
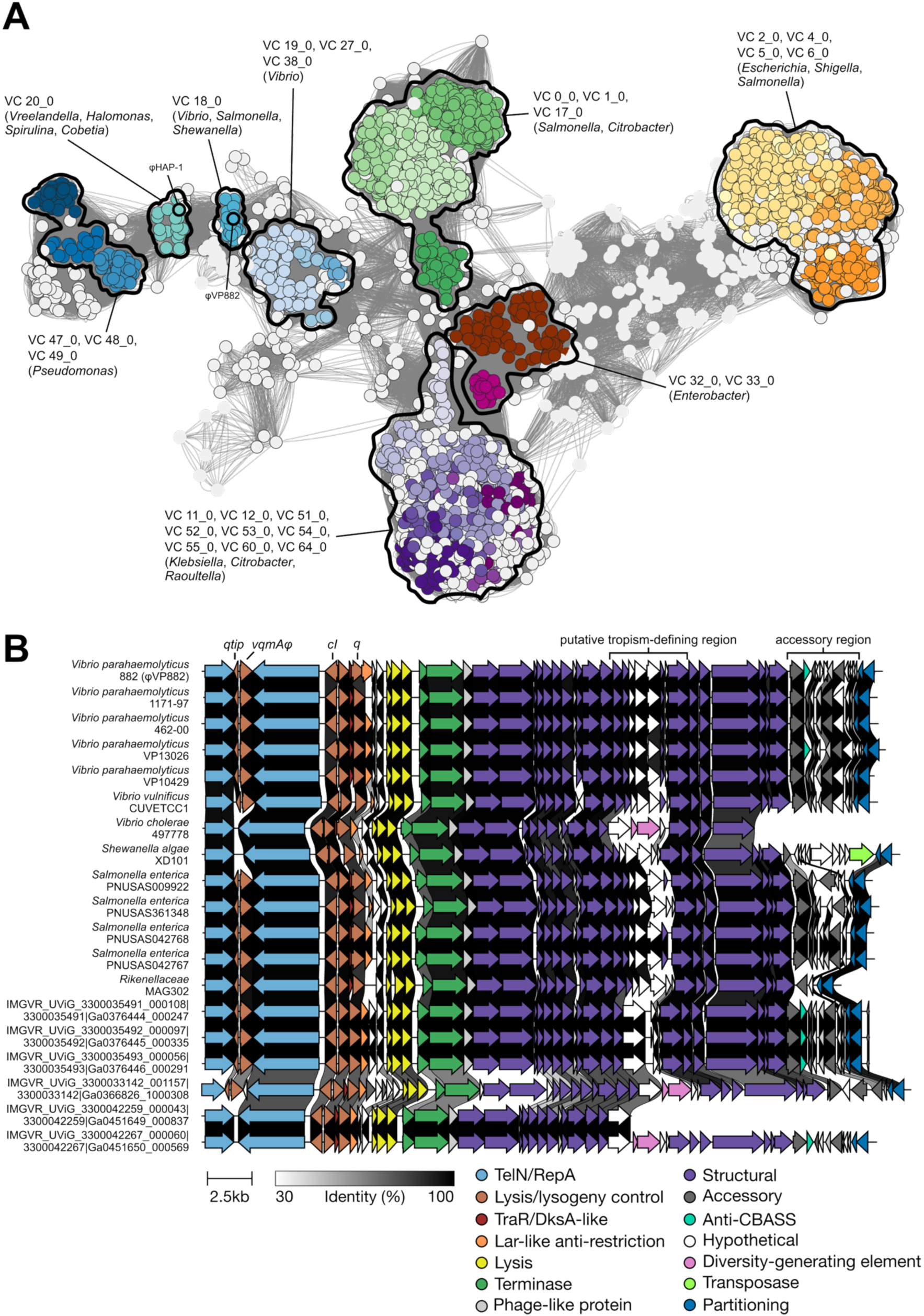
φVP882 clusters with VP882-like linear plasmid phages that lysogenize diverse bacterial species. (A) Taxanomic network of collected linear phage genomes against RefSeq viral sequences. Nodes representing linear phages collected from the 5 queried databases are outlined in black. Nodes from the Prokaryotic Viral RefSeq database are not outlined. Edge weights were calculated by vConTACT2. Viral clusters with more than 20 nodes are colored. Individual clusters or groups of clusters are outlined and labeled with their respective VC numbers and the most abundant associated genera. (B) Genome synteny of VP882-like linear plasmid phages. Host species and strain are provided on the left. Sequences designated IMGVR were collected from metagenomic data and thus, do not have associated host strains. Arrows represent genes colored according to their annotated functions. Gene homologs in neighboring sequences are connected by shaded links. The shading represents the % identity between the amino acid sequences of the proteins encoded by the homologous genes. The absence of a link indicates less than 30% amino acid identity between proteins encoded by neighboring genes or the absence of a homolog in the neighbor.

**Fig 2.**
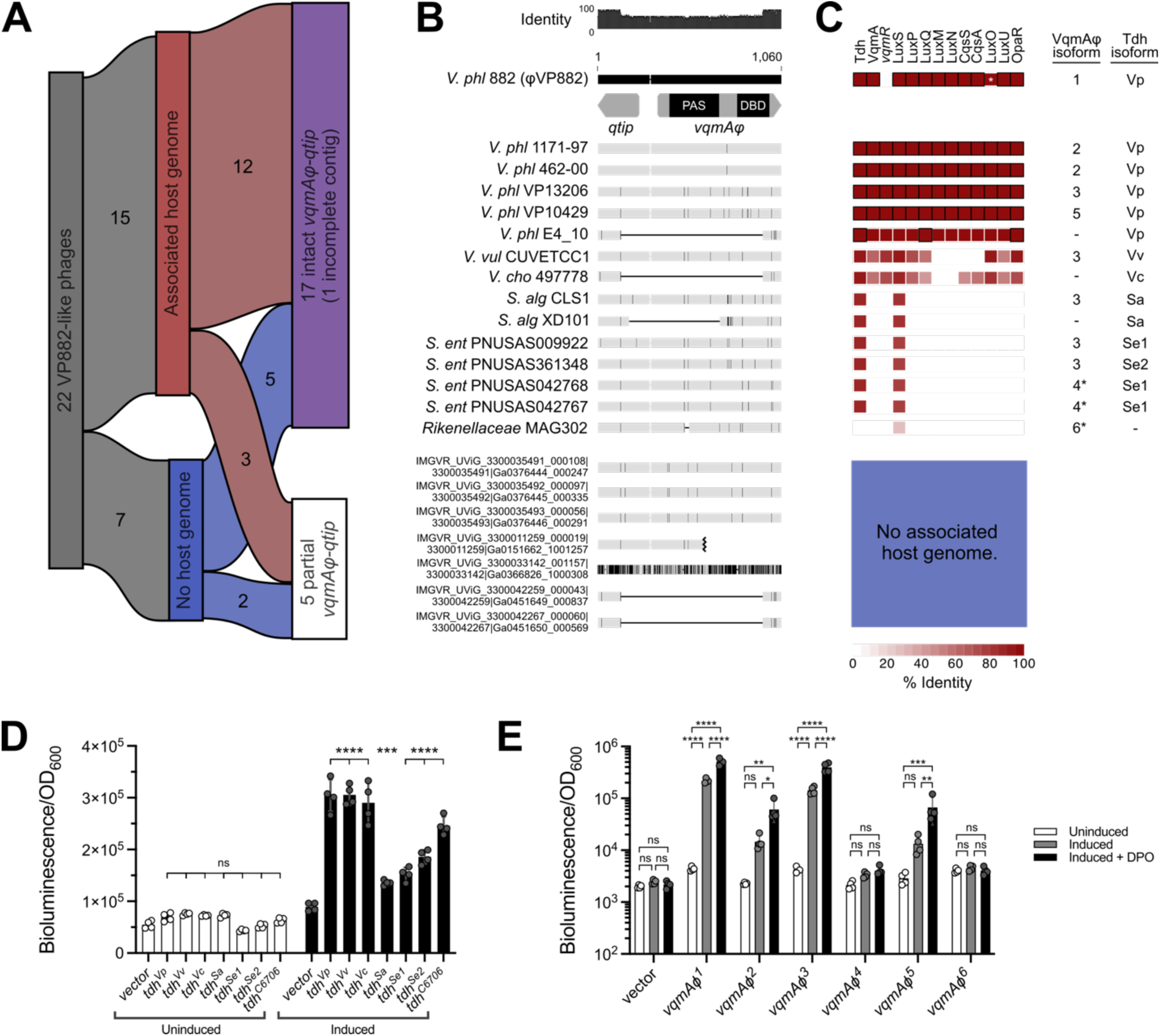
The *vqmA*φ*-qtip* module is conserved and functional across VP882-like linear plasmid phages harbored by diverse DPO-producing bacterial species. (A) Chart showing subsets of VP882-like phages discussed in the text. (B) Nucleotide alignment of the genome region encoding the *qtip* and *vqmA*φ genes from 22 VP882-like phages. A schematic of *qtip* and *vqmA*φ is shown for the uppermost sequence, and the regions encoding the VqmAφ DNA-binding (DBD) and ligand binding (PAS) domains are indicated. The host species and strain are provided on the left (*V. phl* = *Vibrio parahaemolyticus*, *V. vul* = *Vibrio vulnificus*, *V. cho* = *Vibrio cholerae*, *S. alg* = *Shewanella algeae*, *S. ent* = *Salmonella enterica*). Sequences designated IMGVR were collected from metagenomic data and thus, do not have associated host strains. Gray horizontal bars represent the homologous DNA sequences, and black vertical lines within the gray bars represent nucleotide differences relative to *qtip-vqmA*φ from the φVP882 reference sequence. Thin black horizontal lines denote deleted DNA sequences. The black jagged mark in the IMGVR sequence ending with Ga0151662_1001257 denotes the terminus of the contig. (C) Heat map of quorum- sensing protein amino acid identity across host strains carrying VP882-like phages. Identity was calculated relative to quorum-sensing proteins from *V. parahaemolyticus* type strain RIMD 2210633. Black outlines denote 100% identity. Blank spaces indicate no detected homolog. The white asterisk in LuxO from *V. parahaemolyticus* 882 indicates the constitutive low-cell-density- locked status of the protein, despite high homology, due to a 12 amino acid deletion relative to the type strain. Pairs of VqmAφ and Tdh isoforms for each strain are listed to the right of the heatmap. VqmAφ isoforms marked with asterisks are inactive in our assays. (D) Light production from the host-encoded P*_vqmR_-luxCDABE* transcriptional fusion following expression of the *tdh* genes under study in the *V. cholerae* DPO biosensor strain. (E) Light production from a transcriptional reporter of phage-encoded *qtip* (P*_qtip_-luxCDABE*) following expression of the *vqmA*φ genes under study in *V. parahaemolyticus* in the absence and presence of DPO. (D,E) All experiments were performed in biological quadruplicate (n = 4). Symbols represent individual replicate values. Bars represent means. Error bars represent standard deviations. Cultures were treated with water (Untreated) or arabinose and theophylline (Induced). For statistical comparisons, (D) all samples were compared to the vector control within each treatment and (E) all treatments were compared within each sample. Significance was determined by two-way ANOVA with (D) Dunnet’s or (E) Tukey’s multiple comparisons test to determine adjusted p- values: (D) ns = non-significant, ***p = 0.0006, **** p<0.0001, (E) ns = non-significant, *p = 0.0261, **p = 0.0026, 0.0069, ***p = 0.0008, **** p<0.0001.

For the remainder of this study, we focus on the 15 VP882-like phages that have associated host genomes (Fig. 2A-C). Our search yielded 6 species harboring VP882-like phages: *Vibrio parahaemolyticus*, *Vibrio vulnificus*, *Vibrio cholerae*, *Salmonella enterica*, *Shewanella algae*, and a *Rikenellaceae* family isolate (Fig. 1B, Table S1). In *V. parahaemolyticus*, φVP882 lysis-lysogeny transitions are influenced by VqmAφ-Qtip, host DPO production, and the vibrio host LuxO-OpaR quorum-sensing system (13, 14). We determined the presence or absence of vibrio quorum-sensing genes in each host strain harboring a VP882-like phage (Fig. 2C, Fig. S3A-E). Nearly all vibrio quorum-sensing genes are restricted to vibrio hosts. The exceptions are *tdh* (present in 14/15 hosts) and *luxS* (present in 15/15 hosts) (Fig. 2C, Fig. S3A,D).

Tdh and LuxS produce the universal autoinducers DPO and AI-2, respectively. We presume that the VqmAφ-Qtip modules require DPO to drive phage lysis (1). Thus, we focused on Tdh, which is present in each VP882-like phage host strain except *Rikenellaceae* MAG302 (Fig. 2C). Analysis of amino acid identity in Tdh proteins across strains carrying VP882-like phages reveals 6 isoforms that cluster by species (Fig. S3A). To assess whether the Tdh isoforms are functional, we produced each Tdh isoform in a *V. cholerae* Δ*tdh* DPO biosensor strain that carries a transcriptional reporter for *vqmR*, the downstream target of the DPO-VqmA complex. In the reporter, activation of *vqmR* expression tracks linearly with DPO production. All 6 Tdh isoforms produced DPO that activated VqmA in the biosensor strain showing that each Tdh-harboring host strain has the capacity to produce DPO (Fig. 2D). The vibrio Tdh isoforms produced DPO at levels comparable to that from our laboratory *V. cholerae* strain (Tdh^C6706^). Tdh^Sa^, Tdh^Se1^, and Tdh^Se2^ produced less DPO, but we do not know whether the non-vibrio Tdh enzymes are inherently less active than those from vibrios or display sub-optimal function in a recombinant host.

Of the 15 VP882-like phages with associated host genomes, 12 carry an intact *vqmA*φ comprising 6 isoforms (VqmAφ^1-6^) (Fig. 2A-C, Fig. S2A). We assessed whether each of these VqmAφ isoforms, when expressed in a *V. parahaemolyticus* Δ*tdh* strain, could detect exogenously supplied DPO and, in response, activate *qtip* expression. (Fig. 2E). Both apo- and DPO-bound VqmAφ activate *qtip* expression, with holo-VqmAφ being more active (15). VqmAφ^1^, VqmAφ^2^, VqmAφ^3^, and VqmAφ^5^ activated *qtip* expression in the apo-state, and in each case, *qtip* expression increased following DPO supplementation (Fig. 2E). These 4 isoforms are present across 9 of the 12 VP882-like phages with intact *vqmA*φ-*qtip* modules (Fig. 2A,C) and are distributed across *V. parahaemolyticus* (VqmAφ^1,2,3,5^), *V. vulnificus* (VqmAφ^3^), *S. enterica* (VqmAφ^3^), and *S. algae* (VqmAφ^3^) (Fig. 2C, Fig. S2A). VqmAφ^4^ (*S. enterica*) and VqmAφ^6^ (*Rikenellaceae*) failed to activate *qtip* expression in our experimental system (Fig. 2C,E, Fig. S2A). In total, 9 of the 15 VP882-like phages with associated host genomes (Fig. 2A) encode a VqmAφ protein capable of activating the lysis-inducing *qtip* gene in response to host-produced DPO.

## CONCLUSION

We show that φVP882 is a member of a taxon of VP882-like phages that infect diverse bacterial species. All but one of the bacterial hosts encode a functional Tdh, and thus, likely produce DPO. Most VP882-like phages harbor a VqmAφ capable of detecting DPO and, in response, driving *qtip* transcription. Since the characterization of φVP882 as the first temperate phage to integrate host quorum-sensing cues into its lysis-lysogeny decision-making, other variations on quorum-sensing-responsive bacteriophages, including phages that both produce and detect autoinducers, have been identified (1, 16, 17). Our results demonstrate that VP882-like phages, and linear plasmid phages in general, are widely dispersed in nature. By mining expanding viral sequence databases, additional families of quorum-sensing responsive phages that deepen understanding of phage biology and inter-domain chemical communication could be uncovered.

## METHODS

### Viral Taxonomy and Genomic Analysis

Linear plasmid phages were identified using a strategy similar to (17). Searches were performed across NCBI nt, IMG/VR v4 (6), Cenote Human Virome Database v1.1 (7), Global Ocean Virome 2 (5), the Gut Phage Database (8), and the Metagenomic Gut Virus Catalog (9). All sequences were downloaded in April 2025.

*For NCBI nt:* TelN proteins were identified by blastp with the TelN protein sequence from three linear plasmid phages (φVP882 - YP_001039865.1, φ63 – WP_372435025.1, and φ72 - AKN37353.1) (4, 17) against the NCBI nr database. Hits were filtered for >40% identity and >80% query coverage (n=1785). Protein IDs were used to extract phage genomes from NCBI nt. The Identical Protein Groups were gathered for each protein ID and all GenBank contig accession IDs were extracted. These nucleotide accessions were used to fetch 16,409 contigs (length >15kb) and their annotated proteins. Phage status was verified by assessing the collected contigs using VIBRANT (v1.2.1) (18). Sequences called as phages were used in further analyses*. For IMG/VR, CHVD, GOV2, GPD, and MGV*: Database files were annotated with VIBRANT (18) to obtain predicted protein fasta files. TelN proteins were identified by profile HMM search with HMMER3 (v. 3.3.2) (19) for domain PF16684 (Pfam v. 37) (20) against VIBRANT-generated protein databases.

The acquired *telN*-encoding phage genomes were dereplicated with cd-hit-est (21) (parameters: ‘-c 0.99 -aS 1.0 -g 1 -d 0’). All associated protein sequences were deposited into a single combined database and submitted to vConTACT2 (v 0.11.3) (22) with the Prokaryotic Viral RefSeq database (v. 211) (23) to determine linear plasmid phage taxonomy. Genbank files were extracted for all genomes in viral clusters of interest. Two additional VP882-like phages, uncovered by blastp against NCBI nr, were added post-analysis, as their contigs were below the length filter of the analysis pipeline. Whole genome synteny was calculated and visualized with clinker (v. 0.0.31) (24). Genomic alignments were performed in Geneious Prime (v. 2025.0.3) using MUSCLE (25). Heatmaps were generated in R with pheatmap (v. 1.0.13) using % identities calculated in Geneious Prime.

### Host and Viral Protein Functional Assays

Bacterial strains, plasmids, primers, and synthetic DNA fragments used in this study are listed in Tables S2 and S3. Overnight cultures were grown with aeration in Lysogeny Broth (LB-Miller, BD-Difco) at 37° C (*V. cholerae*) or in LB with 2% NaCl (LM) at 30° C (*V. parahaemolyticus*). All experiments were performed in M9 medium with 0.5% glucose, 0.4% casamino acids, and 200 mM NaCl (M9-gluc-CAA-HS). Antibiotics were used at: 50 μg mL^-1^ kanamycin (Kan, GoldBio) and 10 μg mL^-1^ chloramphenicol (Cm, Sigma). L-arabinose (Sigma) and theophylline (Sigma) were used as indicated below.

*For DPO bioassay*: Cultures of the *V. cholerae* DPO biosensor (Δ*tdh* Δ*lacZ*::P*_vqmR_-luxCDABE*) carrying arabinose/theophylline-inducible *tdh* alleles (pXBCm-P*_bad-riboswitch_*-*tdh^X^*) were diluted 1:100 in M9-gluc-CAA-HS supplemented with 10 mM L-threonine (Sigma) and appropriate antibiotics. Tdh production was induced with 0.2% L-arabinose and 200 μM theophylline. *For* qtip *induction assay*: Cultures of *V. parahaemolyticus* (Δ*tdh*) carrying the P*_qtip_*-*luxCDABE* reporter and arabinose/theophylline-inducible *vqmA*φ alleles (pXBCm-P*_bad-riboswitch_*-*vqmA*φ*^X^*) were diluted 1:1000 in M9-gluc-CAA-HS. VqmAφ production was induced with 0.06% arabinose and 200 μM theophylline. 10 μM DPO or water (vehicle) was administered to uninduced and induced samples.

*For both assays*: Diluted cultures were dispensed (200 μL) into white-wall/clear-bottom 96-well plates (Corning Costar). Optical density and bioluminescence were measured with a BioTek Synergy Neo2 Multi-Mode plate reader.

## DATA AVAILABILITY

All datasets used in this study are publicly available. Custom Python scripts used in this study are available upon request.

## ACKNOWLEDGEMENTS

We are grateful to members of the Bassler Laboratory for insightful discussions.

The authors acknowledge financial support from the Howard Hughes Medical Institute, the National Institutes of Health grant R37GM065859, and the National Science Foundation grant MCB-2508324 (B.L.B.).

## SUPPLEMENTARY MATERIAL

**Fig S1.**
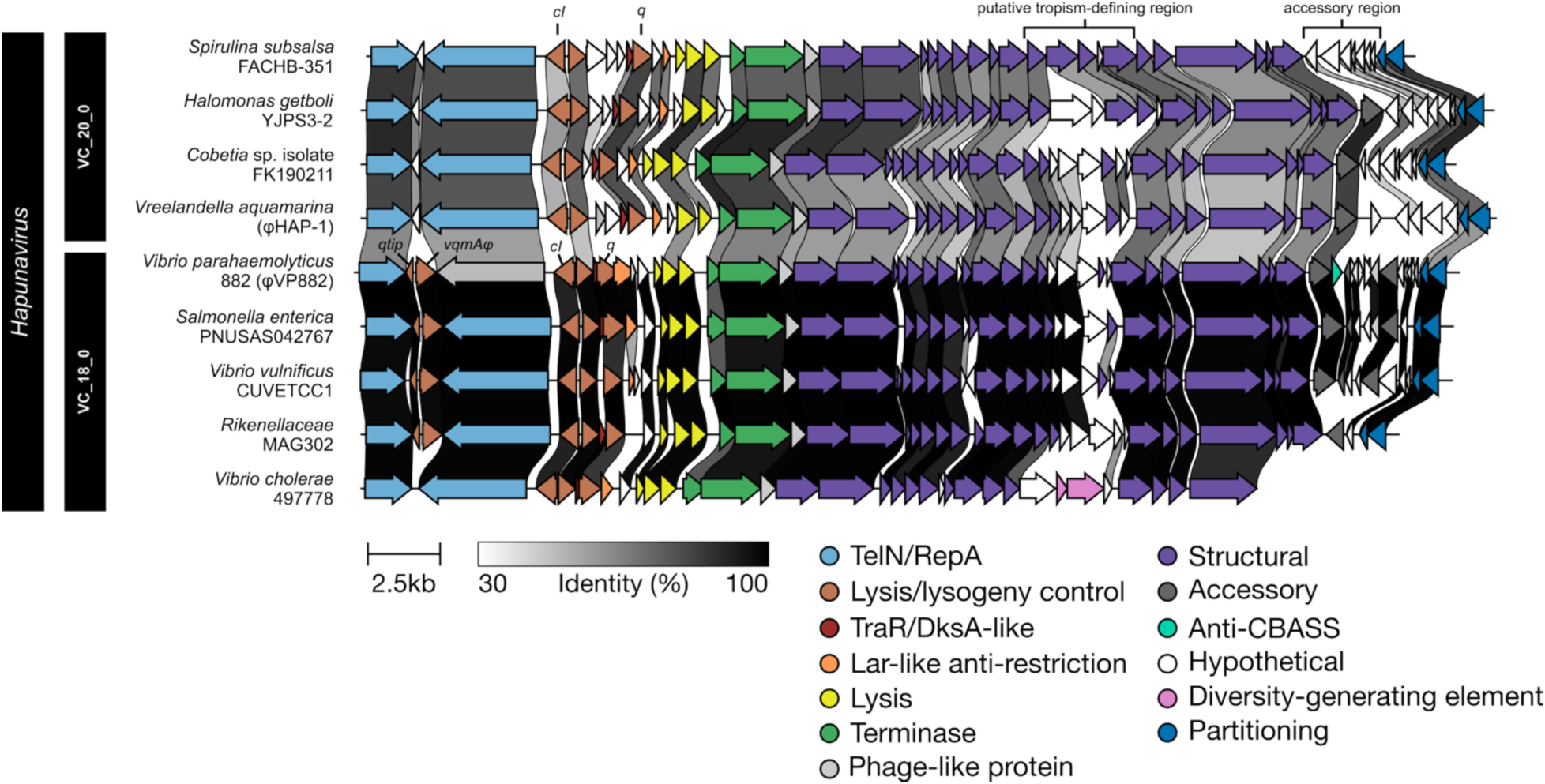
Hapunaviruses are subdivided into VP882-like phages and HAP-1-like phages. Genome synteny of HAP-1-like (VC_20_0) and VP882-like (VC_18_0) linear plasmid phages. Host species and strain are provided on the left. Arrows represent genes colored according to their annotated functions. Gene homologs in neighboring sequences are connected by shaded links. The shading represents the % identity between the amino acid sequences of the proteins encoded by the homologous genes. The absence of a link indicates less than 30% amino acid identity between proteins encoded by neighboring genes or the absence of a homolog in the neighbor.

**Fig S2.**
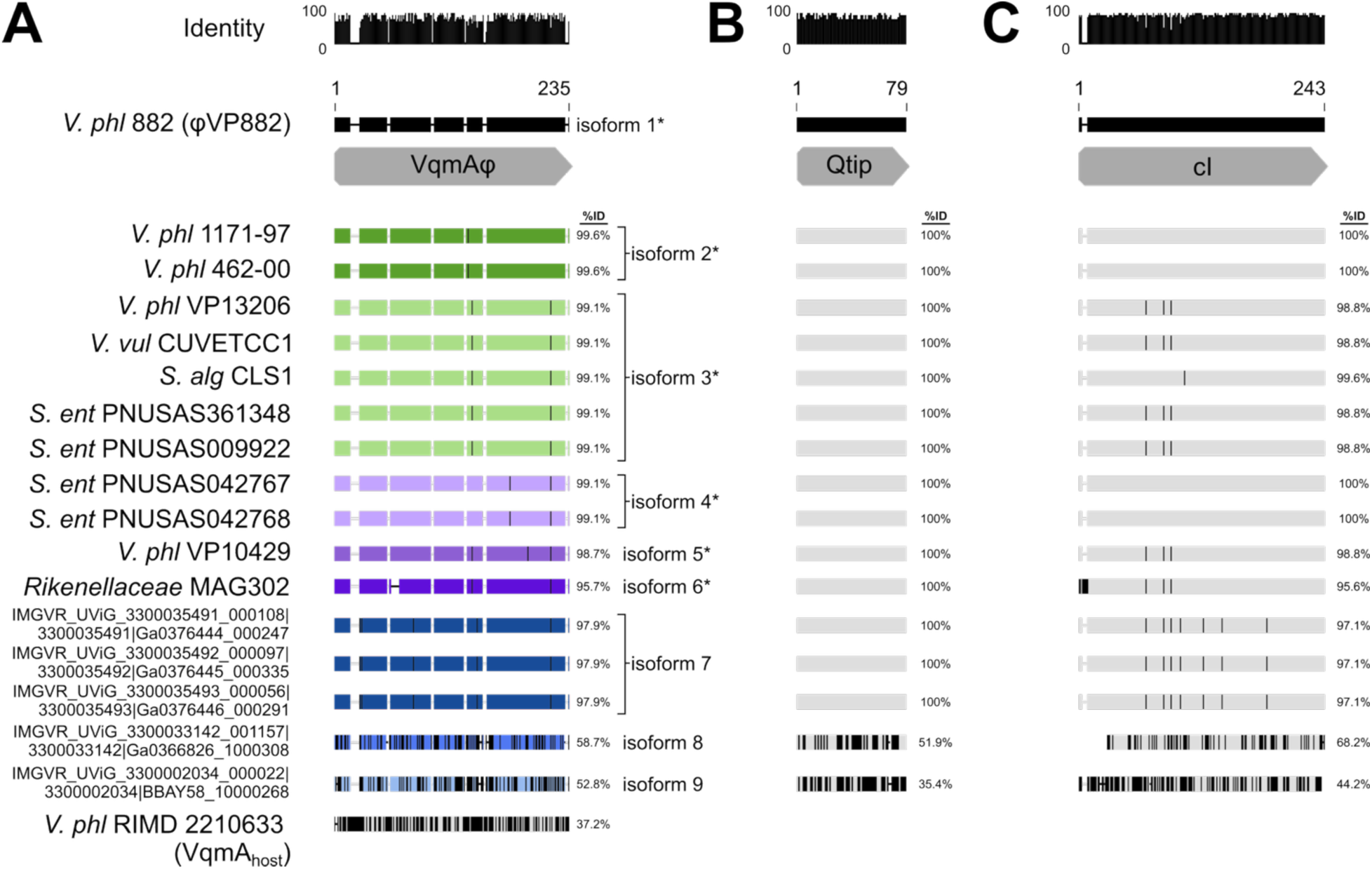
VqmAφ and cI vary across VP882-like phages, whereas Qtip is highly conserved. Pairwise amino acid alignments of (A) VqmAφ, (B) Qtip, and (C) cI proteins from the 17 VP882-like phages with intact *vqmA*φ*-qtip* modules (see Fig. 2A of the main text). Host species and strain are provided on the left (*V. phl* = *Vibrio parahaemolyticus*, *V. vul* = *Vibrio vulnificus*, *V. cho* = *Vibrio cholerae*, *S. ent* = *Salmonella enterica*, *S. alg* = *Shewanella algeae*). Sequences designated IMGVR were collected from metagenomic data and thus, do not have associated host strains. Horizontal bars represent the homologous protein sequences, and black vertical lines within the bars represent amino acid differences relative to the VqmAφ, Qtip, and cI proteins from φVP882. Thin black horizontal lines denote gaps in sequences compared to the reference sequence. Thin gray horizontal lines denote gaps in sequences compared to any sequence other than the reference sequence. In (A), VqmAφ proteins are colored and labeled by isoform. Isoforms marked with asterisks were used in assays in the main text. The VqmA_host_ protein sequence from *V. parahaemolyticus* str RIMD 2210633 was included to demonstrate that while VqmAφ isoforms 8 and 9 vary most among phage VqmAφ homologs, they share greater identity with VqmAφ than the identity shared between VqmA_host_ and VqmAφ. This finding supports the logic that all the VqmAφ isoforms shown should be considered VqmAφ rather than some other transcription factor.

**Fig S3.**
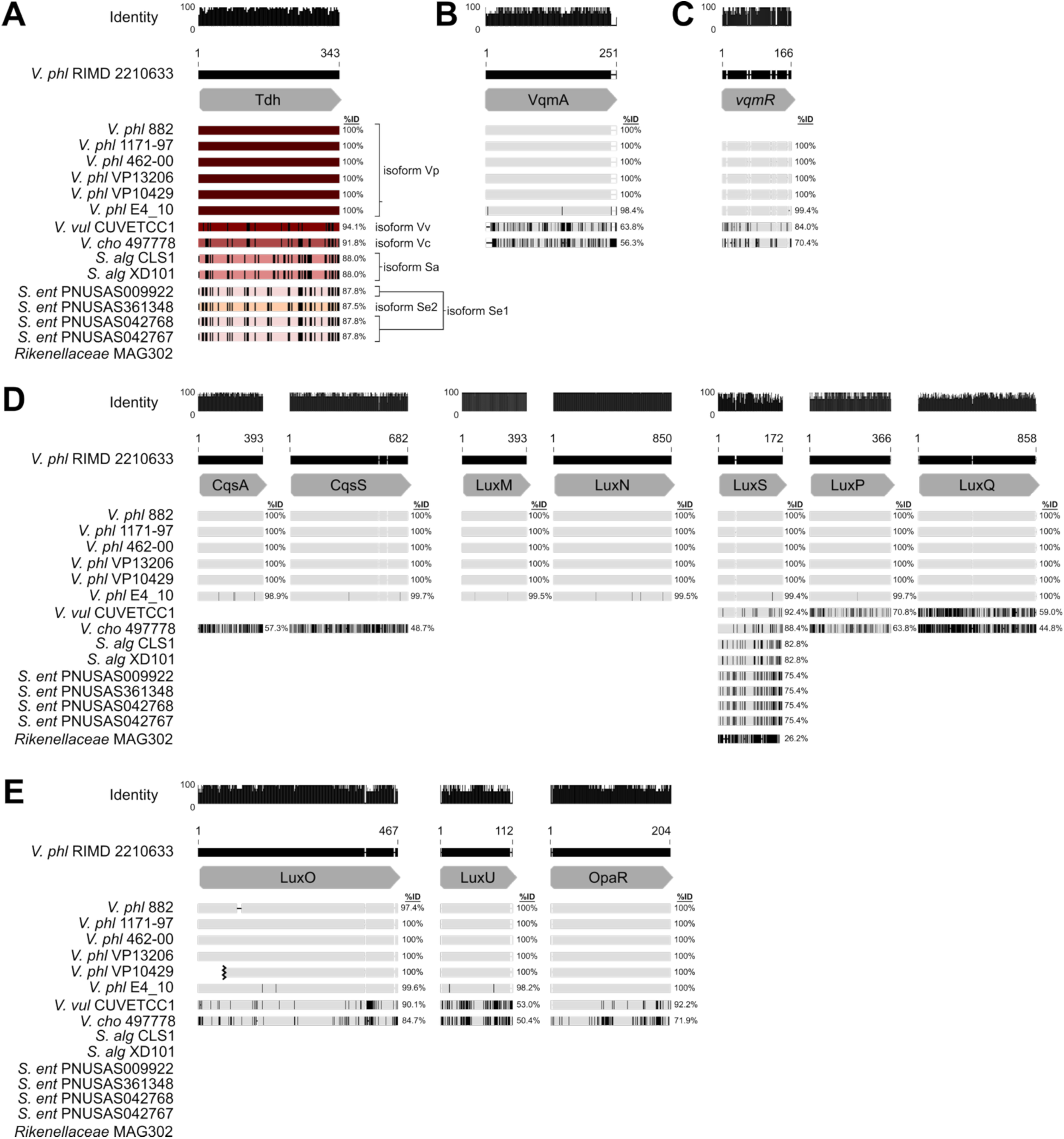
LuxO-OpaR(LuxR) and VqmAR quorum-sensing proteins are restricted to vibrios, while the autoinducer synthases Tdh and LuxS are conserved across genera. Pairwise alignments of (A) Tdh, (B) VqmA_host_, (C) *vqmR*, (D) the autoinducer synthase and receptor proteins associated with the LuxO-OpaR pathway, and (E) LuxO, LuxU, and OpaR from 15 host strains harboring VP882-like phages (see Fig. 2A of the main text). (A-E) Host species and strain are provided on the left (*V. phl* = *Vibrio parahaemolyticus*, *V. vul* = *Vibrio vulnificus*, *V. cho* = *Vibrio cholerae*, *S. ent* = *Salmonella enterica*, *S. alg* = *Shewanella algeae*). Horizontal bars represent the homologous (A,B,D,E) protein or (C) nucleotide sequences, and black vertical lines within the bars represent (A,B,D,E) amino acid or (C) nucleotide differences relative to the corresponding sequences from the *V. parahaemolyticus* reference strain RIMD 2210633. Thin black horizontal lines denote gaps in sequences compared to the reference sequence. Thin gray horizontal lines denote gaps in sequences compared to any sequence other than the reference sequence. (A) Tdh proteins colored and labeled by isoform. (E) The black jagged mark in the *V. phl* VP10429 sequence denotes the end of the contig.

**Table S1.** Genome metadata for VP882-like linear plasmid phages.

| Host Strain | vConTACT2<br>Viral Cluster | Isolation<br>Date | Isolation<br>Location | Source |
| --- | --- | --- | --- | --- |
| *Vibrio parahaemolyticus* 882 (1) | VC_18_0 | unknown<br>(published<br>2009) | Taiwan | unknown |
| *Vibrio parahaemolyticus* 1171-97 | VC_18_0 | 1997 | Peru: Moquegua | clinical |
| *Vibrio parahaemolyticus* 462-00 | VC_18_0 | 2000 | Peru: Lima | clinical |
| * *Vibrio parahaemolyticus* 461-00 | n/a | 2000 | Peru: Lima | clinical |
| ** *Vibrio parahaemolyticus* E4_10 | n/a | 2014 | China: Zhejiang | fish |
| *Vibrio parahaemolyticus* str VP13026 | VC_18_0 | 2013 | China: Shenzhen | clinical |
| *Vibrio parahaemolyticus* str VP10429 | VC_18_0 | 2010 | China: Shenzhen | clinical |
| *Vibrio vulnificus* str CUVETCC1 | VC_18_0 | 2021 | Thailand:<br>Chachoengsao | fish |
| *Vibrio cholerae* str 497778 | VC_18_0 | 2017 | unknown | clinical |
| *Salmonella enterica* str<br>PNUSAS042767 | VC_18_0 | 2018 | USA | clinical |
| *Salmonella enterica* str<br>PNUSAS042768 | VC_18_0 | 2018 | USA | clinical |
| *Salmonella enterica* str<br>PNUSAS361348 | VC_18_0 | 2023 | USA | clinical |
| *Salmonella enterica* str<br>PNUSAS009922 | VC_18_0 | 2017 | USA | clinical |
| *Shewanella algae* str XD101 | VC_18_0 | 2023 | China: Sanya | seawater |
| ** *Shewanella algae* str CLS1 | n/a | 2014 | Taiwan | clinical |
| *Rikenellaceae* bacterium MAG302 | VC_18_0 | 2017 | New Zealand:<br>Little Barrier<br>Island, Auckland | fish gut<br>metagenome |
| IMGVR_UViG_3300011259_000019,<br>3300011259, Ga0151662_1001257 | VC_18_0 | 2015 | Japan: Japan<br>Sea near Toyama<br>Prefecture | marine<br>sediment |
| IMGVR_UViG_3300035491_000108,<br>3300035491, Ga0376444_000247 | VC_18_0 | 2007 | Trinidad and<br>Tobago: La Brea,<br>Pitch Lake | asphalt lake |
| IMGVR_UViG_3300035492_000097,<br>3300035492, Ga0376445_000335 | VC_18_0 | 2007 | Trinidad and<br>Tobago: La Brea,<br>Pitch Lake | asphalt lake |
| *IMGVR_UViG_3300035494_000080,<br>3300035494, Ga0376447_000328 | n/a | 2007 | Trinidad and<br>Tobago: La Brea,<br>Pitch Lake | asphalt lake |
| IMGVR_UViG_3300035493_000056,<br>3300035493, Ga0376446_000291 | VC_18_0 | 2007 | Trinidad and<br>Tobago: La Brea,<br>Pitch Lake | asphalt lake |
| IMGVR_UViG_3300042259_000043,<br>3300042259, Ga0451649_000837 | VC_18_0 | 2019 | USA: Denver,<br>Colorado | industrial<br>wastewater |
| IMGVR_UViG_3300042267_000060,<br>3300042267, Ga0451650_000569 | VC_18_0 | 2019 | USA: Denver,<br>Colorado | industrial<br>wastewater |
| *IMGVR_UViG_3300042269_000097,<br>3300042269, Ga0451652_000962 | n/a | 2019 | USA: Denver,<br>Colorado | industrial<br>wastewater |
| IMGVR_UViG_3300033142_001157,<br>3300033142, Ga0366826_1000308 | VC_18_0 | unknown<br>(added to<br>database<br>2019) | Mexico: Gulf of<br>California | marine<br>sediment |
| †IMGVR_UViG_3300002034_000022,<br>3300002034, BBAY58_10000268 | Outlier | unknown<br>(added to<br>database<br>2013) | Australia: Sydney,<br>Bare Island | red algae |

\* Viral contig excluded from vConTACT2 due to >99% similarity to the entry directly above it in the table.
\*\* Viral contigs identified by blastp but excluded from phage search due to contig lengths.
† Viral contig encodes VqmAϕ-Qtip but does not cluster with VP882-like phages.

**Table S2.** Strains and plasmids used in this study.

| **Bacterial Strains** |  |  |  |
| --- | --- | --- | --- |
| **Parent Strain** | **Identifier** | **Genotype** | **Reference** |
| *V. cholerae* C6706 | BB-Vc0325 | $\Delta tdh \Delta lacZ::P_{vqmR}-luxCDABE$ | (2) |
| BB-Vc0325 | FJS-S1644 | /pXBCm-P<sub>bad-riboswitch</sub>-gfp | This study |
|  | FJS-S1645 | /pXBCm-P<sub>bad-riboswitch</sub>-tdh<sup>Vp</sup> | This study |
|  | FJS-S1648 | /pXBCm-P<sub>bad-riboswitch</sub>-tdh<sup>Vv</sup> | This study |
|  | FJS-S1649 | /pXBCm-P<sub>bad-riboswitch</sub>-tdh<sup>Vc</sup> | This study |
|  | FJS-S1650 | /pXBCm-P<sub>bad-riboswitch</sub>-tdh<sup>Sa</sup> | This study |
|  | FJS-S1651 | /pXBCm-P<sub>bad-riboswitch</sub>-tdh<sup>Se1</sup> | This study |
|  | FJS-S1652 | /pXBCm-P<sub>bad-riboswitch</sub>-tdh<sup>Se2</sup> | This study |
|  | FJS-S1653 | /pXBCm-P<sub>bad-riboswitch</sub>-tdh<sup>C6706</sup> | This study |
| *V. parahaemolyticus* RIMD 2210633 | FJS-S0063 | $\Delta tdh$; Amp<sup>R</sup>, Sm<sup>R</sup> | This study |
| FJS-S0063 | FJS-S1614 | /pEVS143-P<sub>qtip</sub>-luxCDABE<br>/pXBCm-P<sub>bad-riboswitch</sub>-gfp | This study |
| | FJS-S1615 | /pEVS143-P<sub>qtip</sub>-luxCDABE<br>/pXBCm-P<sub>bad-riboswitch</sub>-vqmA$\phi^1$ | This study |
| | FJS-S1655 | /pEVS143-P<sub>qtip</sub>-luxCDABE<br>/pXBCm-P<sub>bad-riboswitch</sub>-vqmA$\phi^2$ | This study |
| | FJS-S1656 | /pEVS143-P<sub>qtip</sub>-luxCDABE<br>/pXBCm-P<sub>bad-riboswitch</sub>-vqmA$\phi^3$ | This study |
| | FJS-S1657 | /pEVS143-P<sub>qtip</sub>-luxCDABE<br>/pXBCm-P<sub>bad-riboswitch</sub>-vqmA$\phi^4$ | This study |
| | FJS-S1658 | /pEVS143-P<sub>qtip</sub>-luxCDABE<br>/pXBCm-P<sub>bad-riboswitch</sub>-vqmA$\phi^5$ | This study |
| | FJS-S1659 | /pEVS143-P<sub>qtip</sub>-luxCDABE<br>/pXBCm-P<sub>bad-riboswitch</sub>-vqmA$\phi^6$ | This study |
| *E. coli* TOP10 | | F- mcrA $\Delta(mrr-hsdRMS-mcrBC)$ $\phi 80lacZ\Delta M15$ $\Delta lacX74$ recA1 araD139 $\Delta(ara-leu)$ 7697 galU galK λ- rpsL(Str<sup>R</sup>) endA1 nupG | Invitrogen |
| **Plasmids** |  |  |  |
| **Name** | **Identifier** | **Description** | **Reference** |
| pXBCm-P<sub>bad-riboswitch</sub>-gfp | FJS-P122 | Dual-control expression construct (arabinose-inducible transcription/theophylline-inducible translation) on a chloramphenicol-resistant version of the pXB300 (3) backbone; ColE1; Cm<sup>R</sup> | This study |
| pXBCm-P<sub>bad-riboswitch</sub>-tdh<sup>Vp</sup> | FJS-P127 | *Vibrio parahaemolyticus* 882 *tdh* allele expression plasmid; ColE1; Cm<sup>R</sup> | This study |
| pXBCm-P<sub>bad-riboswitch-</sub><br>*tdh*<sup>Vv</sup> | FJS-P131 | *Vibrio vulnificus* CUVETCC1 *tdh* allele expression plasmid; ColE1; Cm<sup>R</sup> | This study |
| pXBCm-P<sub>bad-riboswitch-</sub><br>*tdh*<sup>Vc</sup> | FJS-P132 | *Vibrio cholerae* 497778 *tdh* allele expression plasmid; ColE1; Cm<sup>R</sup> | This study |
| pXBCm-P<sub>bad-riboswitch-</sub><br>*tdh*<sup>Sa</sup> | FJS-P133 | *Shewanella algae* CLS1 *tdh* allele expression plasmid; ColE1; Cm<sup>R</sup> | This study |
| pXBCm-P<sub>bad-riboswitch-</sub><br>*tdh*<sup>Se1</sup> | FJS-P134 | *Salmonella enterica* PNUSAS042767 *tdh* allele expression plasmid; ColE1; Cm<sup>R</sup> | This study |
| pXBCm-P<sub>bad-riboswitch-</sub><br>*tdh*<sup>Se2</sup> | FJS-P135 | *Salmonella enterica* PNUSAS361348 *tdh* allele expression plasmid; ColE1; Cm<sup>R</sup> | This study |
| pXBCm-P<sub>bad-riboswitch-</sub><br>*tdh*<sup>C6706</sup> | FJS-P136 | *Vibrio cholerae* C6706 *tdh* allele expression plasmid; ColE1; Cm<sup>R</sup> | This study |
| pEVS143-P<sub>qtip-</sub><br>*luxCDABE* | pOD-58 | Transcriptional fusion of the ϕVP882 *qtip* promoter to the *luxCDABE* operon; p15A; Km<sup>R</sup> | (4) |
| pXBCm-P<sub>bad-riboswitch-</sub><br>*vqmA*ϕ<sup>1</sup> | FJS-P128 | *Vibrio parahaemolyticus* 882 *vqmA*ϕ allele expression plasmid; ColE1; Cm<sup>R</sup> | This study |
| pXBCm-P<sub>bad-riboswitch-</sub><br>*vqmA*ϕ<sup>2</sup> | FJS-P138 | *Vibrio parahaemolyticus* 1171-97 *vqmA*ϕ allele expression plasmid; ColE1; Cm<sup>R</sup> | This study |
| pXBCm-P<sub>bad-riboswitch-</sub><br>*vqmA*ϕ<sup>3</sup> | FJS-P139 | *Vibrio parahaemolyticus* VP13206 *vqmA*ϕ allele expression plasmid; ColE1; Cm<sup>R</sup> | This study |
| pXBCm-P<sub>bad-riboswitch-</sub><br>*vqmA*ϕ<sup>4</sup> | FJS-P140 | *Salmonella enterica* PNUSAS042767 *vqmA*ϕ allele expression plasmid; ColE1; Cm<sup>R</sup> | This study |
| pXBCm-P<sub>bad-riboswitch-</sub><br>*vqmA*ϕ<sup>5</sup> | FJS-P141 | *Vibrio parahaemolyticus* VP10429 *vqmA*ϕ allele expression plasmid; ColE1; Cm<sup>R</sup> | This study |
| pXBCm-P<sub>bad-riboswitch-</sub><br>*vqmA*ϕ<sup>6</sup> | FJS-P142 | *Rikenellaceae* bacterium MAG302 *vqmA*ϕ allele expression plasmid; ColE1; Cm<sup>R</sup> | This study |

**Table S3.**
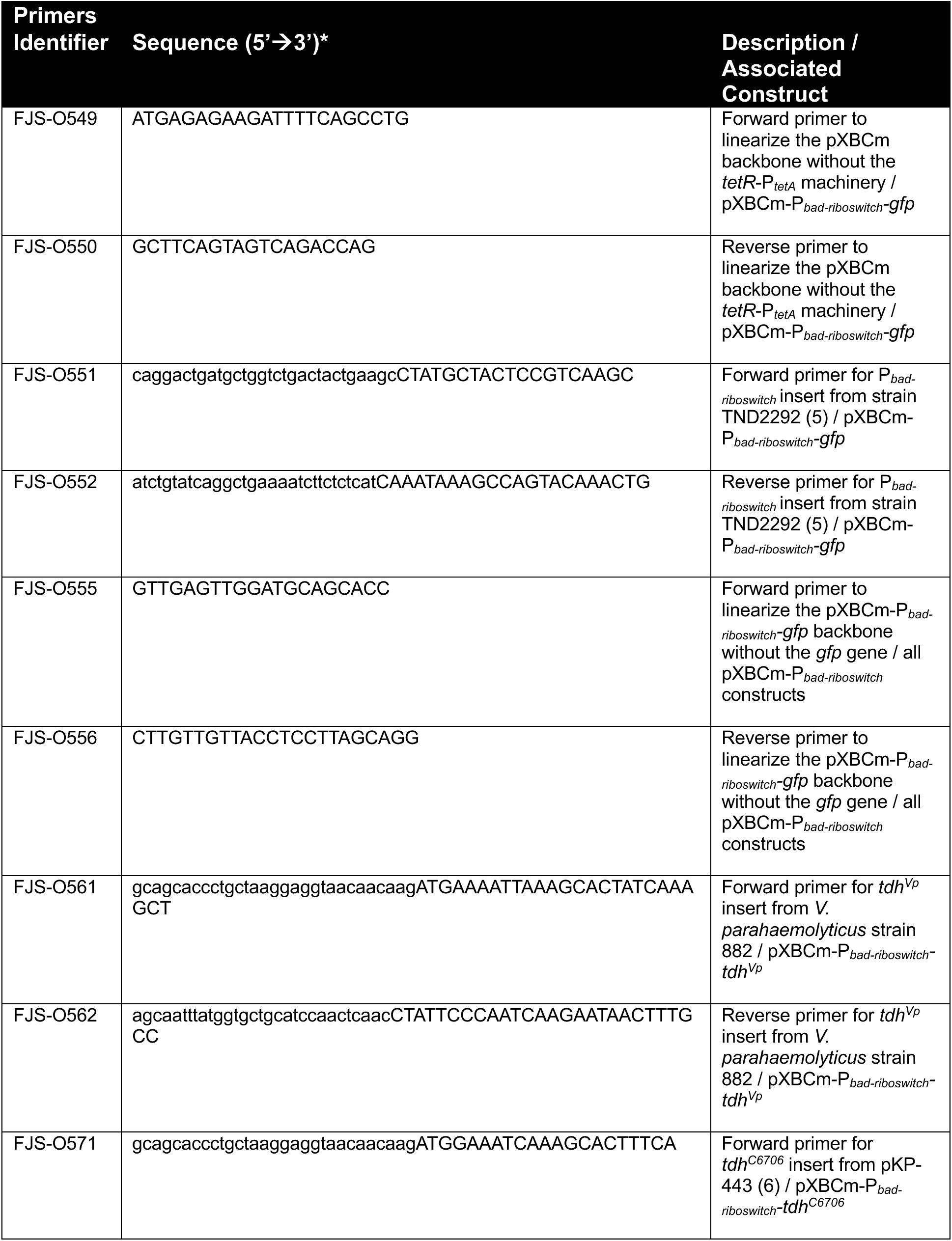
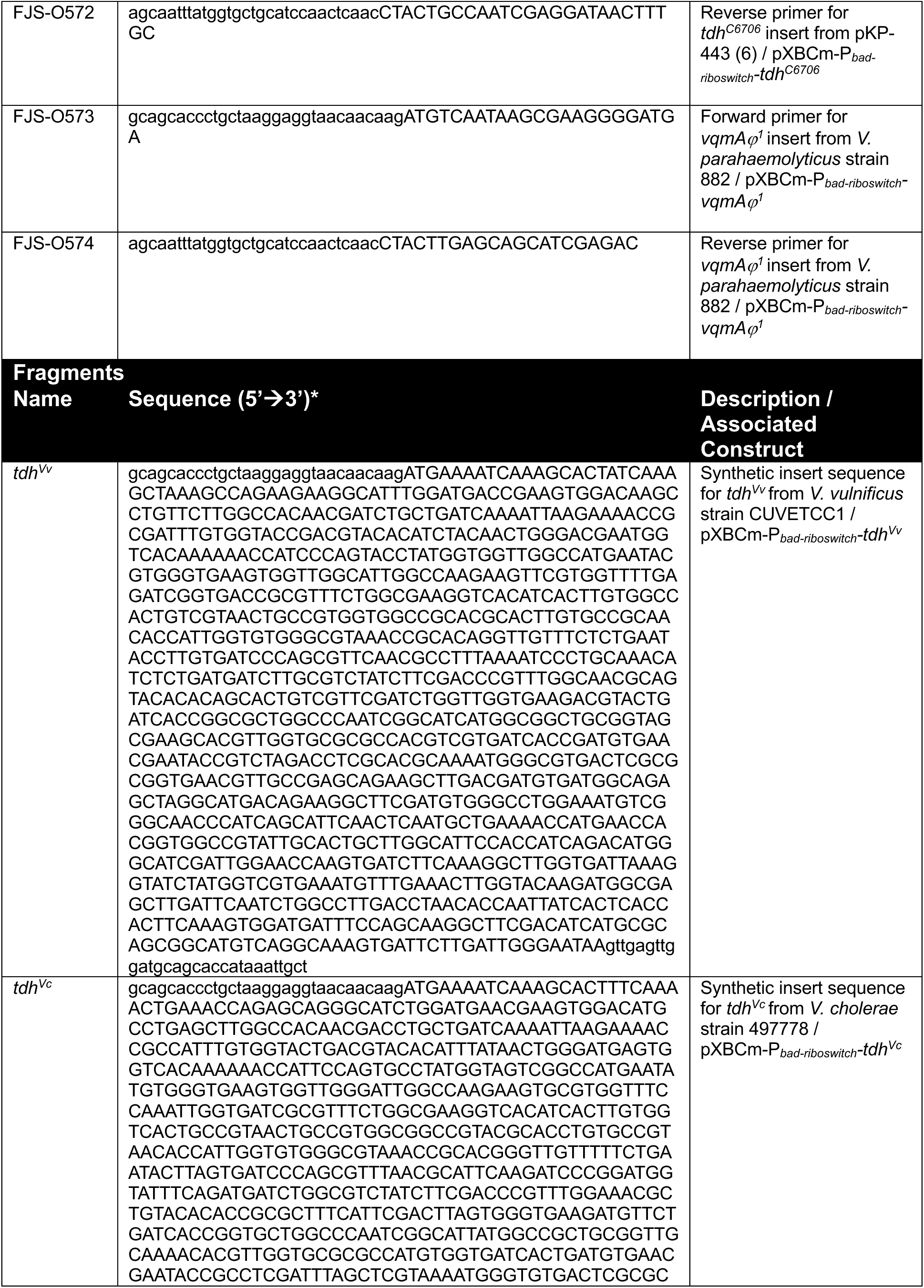
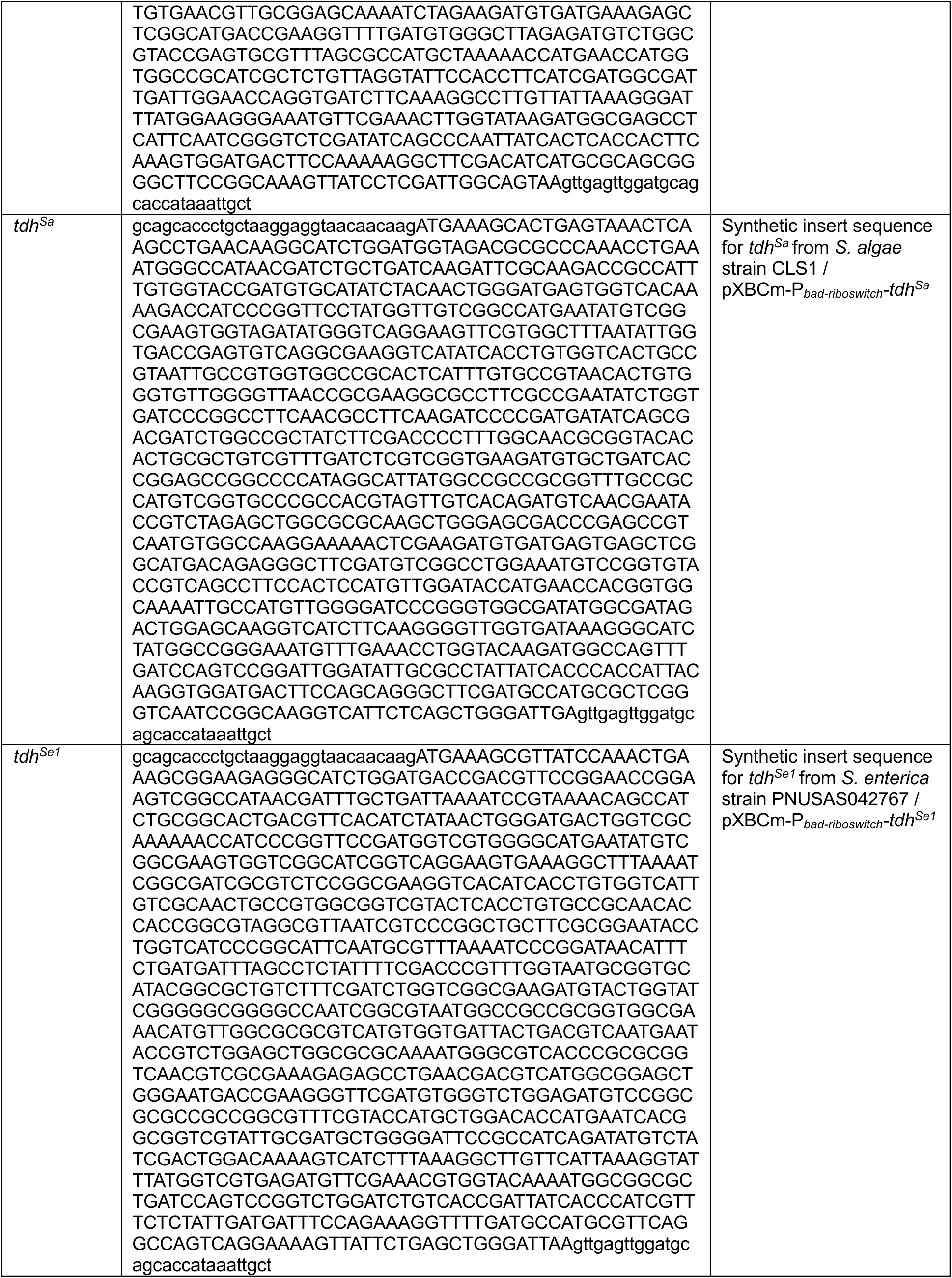
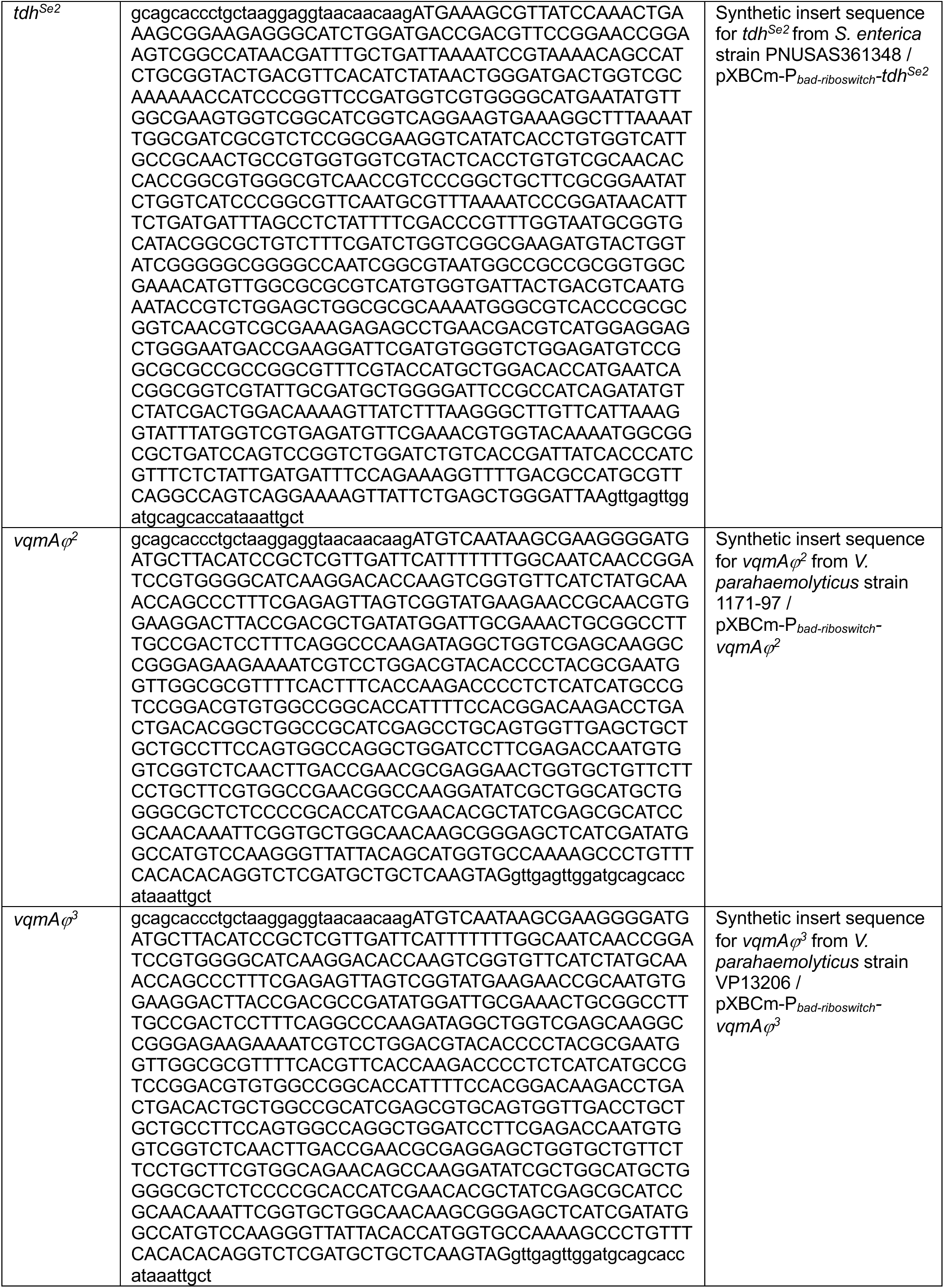
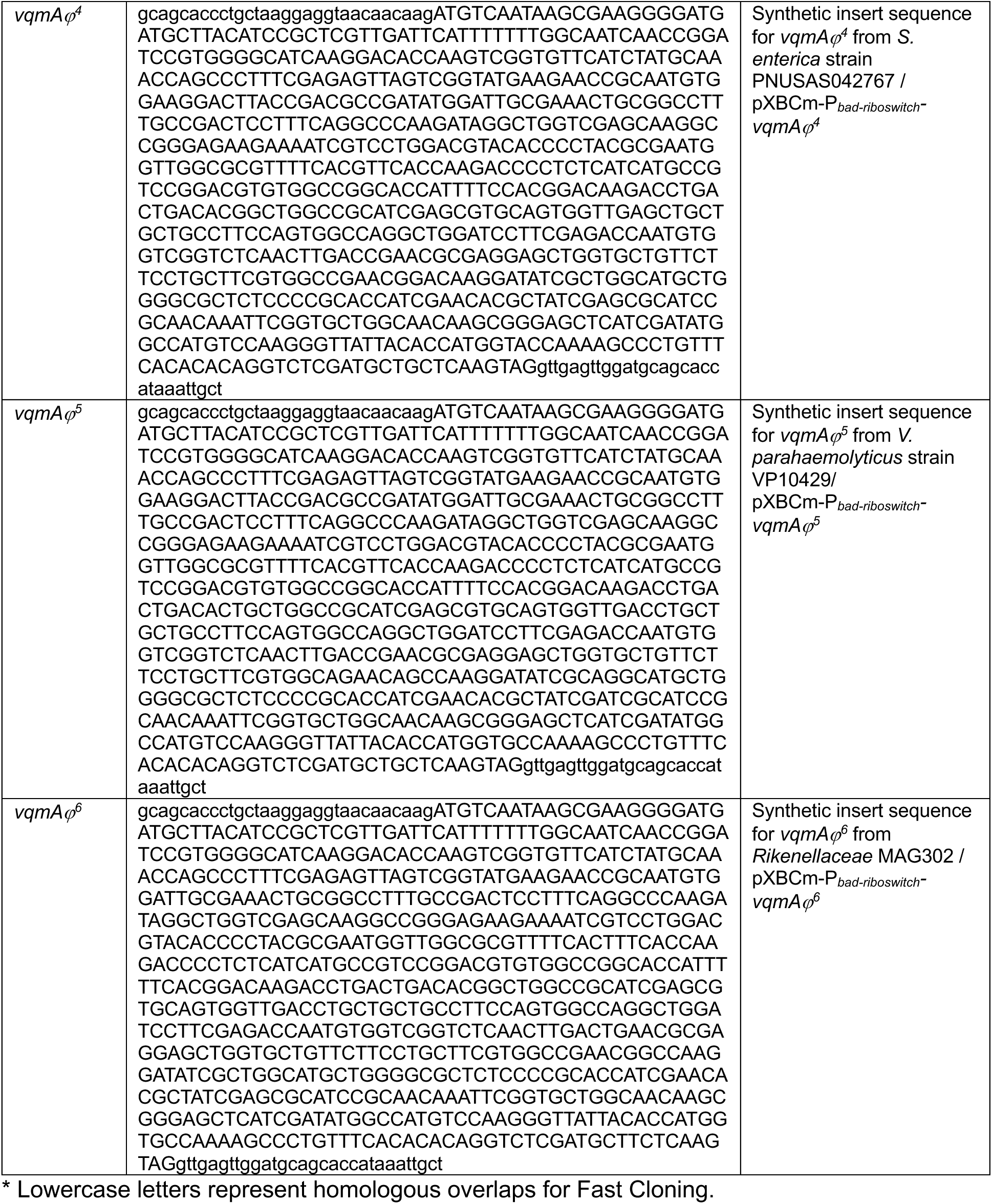
Primers and synthetic DNA fragments used in this study.

**Supplementary Dataset 1.** Identifiers for all genomes used in this study.

**Supplementary Dataset 2.** vConTACT assigned viral clusters for all phage genomes.

